# Susceptibility to Task-irrelevant Auditory Distractors in Relation to Visual Working Memory in Children With and Without ADHD

**DOI:** 10.1101/2025.11.27.691069

**Authors:** Yuanjun Kong, Manqi Zhou, Xuye Yuan, Xiangsheng Luo, Jipeng Huang, Yiwen Li, Lili Yang, Yiyang Wang, Li Sun, Yan Song

**Affiliations:** Key Laboratory of Adolescent Cyberpsychology and Behavior (CCNU), Ministry of Education, Wuhan 430079, China; Key Laboratory of Human Development and Mental Health of Hubei Province, School of Psychology, Central China Normal University, Wuhan 430079, China; State Key Laboratory of Cognitive Neuroscience and Learning, Beijing Normal University, Beijing 100875, China; The National Clinical Research Center for Mental Disorders & Beijing Key Laboratory of Mental Disorders, Beijing Anding Hospital, Capital Medical University, Beijing 100088, China; Faculty of Education, Beijing Normal University, Beijing 100875, China; Peking University Sixth Hospital & Peking University Institute of Mental Health, Beijing 100191, China; NHC Key Laboratory of Mental Health (Peking University) & National Clinical Research Center for Mental Disorders (Peking University Sixth Hospital), Beijing 100191, China

**Keywords:** ADHD, MMN, visual working memory, auditory distraction, EEG

## Abstract

**Objective:** Attention-deficit/hyperactivity disorder (ADHD) is a prevalent neurodevelopmental disorder in children and is often associated with increased distractibility. Previous research has reported that susceptibility to task-irrelevant visual distractions is associated with visual working memory (WM) capacity. However, how neural responses to task-irrelevant auditory distractors are related to visual WM capacity in children with and without ADHD remains poorly understood.

**Methods:** In this study, we collected electroencephalography (EEG) signals from 72 children with ADHD and 88 typically developing (TD) children across two experiments, during which they performed visual tasks of varying difficulty while task-irrelevant auditory distractors were presented. The mismatch negativity (MMN) amplitude was measured as a neural index of susceptibility to auditory distractors, and the K score was assessed as a behavioral indicator of visual WM capacity.

**Results:** TD children showed opposite patterns across the two experiments: under low audiovisual competition, those with higher visual working memory capacity were more susceptible to task-irrelevant sounds, whereas under high competition, children with lower capacity showed greater susceptibility to distraction. In contrast, children with ADHD showed a stable association, with lower visual WM capacity consistently predicting higher susceptibility to task-irrelevant auditory distractors regardless of competition level.

**Conclusions:** These findings provide novel evidence that susceptibility to task-irrelevant auditory distraction is closely related to visual WM capacity in both children with and without ADHD, but the patterns of this relationship differ between typical and atypical neurodevelopment and are modulated by audiovisual competition.

## Introduction

Attention-deficit/hyperactivity disorder (ADHD) is a common neurodevelopmental disorder in childhood characterized by inappropriate patterns of inattentiveness, hyperactivity and/or impulsivity (American Psychiatric Association, 2013), affecting approximately 7.2% of children globally (Thomas, Sanders, Doust, Beller, & Glasziou, 2015). These core symptoms are often accompanied by impairments in executive functions, particularly working memory (WM), which is critical for regulating behavior and cognition (Biederman et al., 2006).

The increased distractibility observed in children with ADHD is hypothesized to reflect a limited capacity to filter or suppress irrelevant stimuli, indicating impairments in attentional control (Mason, Humphreys, & Kent, 2005) or filtering efficiency (Jonkman, Kenemans, Kemner, Verbaten, & Van Engeland, 2004). The attentional filtering mechanism has been thought to be related to individual differences in WM capacity (Burgoyne & Engle, 2020). In adults, individuals with lower visual WM capacity show less efficient attentional filtering, allowing irrelevant information to consume limited memory resources, whereas those with higher capacity can better suppress distractors and maintain relevant input (McNab & Klingberg, 2008). This view is supported by event-related potential (ERP) findings from (Gaspar, Christie, Prime, Jolicœur, & McDonald, 2016), who reported that adults with low visual WM capacity were more susceptible to attentional capture by task-irrelevant visual distractors, whereas those with high capacity could prevent such capture through suppression. Consistent patterns have been observed in behavioral studies in children with and without ADHD. For example, more efficient filtering of task-irrelevant distractors was associated with greater WM performance in typically developing (TD) children (Plebanek & Sloutsky, 2019), and inattentive behaviors, such as looking toward irrelevant interference, were related to impaired WM storage in children with ADHD (Kofler, Rapport, Bolden, Sarver, & Raiker, 2010). However, recent ERP evidence from Zhong et al. (2024) suggests that this relation may be reversed when task-relevant distractors contain target-defining features and are presented sequentially with targets. Adults with higher visual WM capacity showed greater contingent attentional capture by task-relevant distractors, indicating that the relationship between visual WM and distractibility may be context-dependent.

While most research has focused on the relationship between visual WM capacity and susceptibility to visual distractors, few studies have investigated how auditory distraction relates to visual WM capacity. In the limited behavioral evidence, adults and TD children with higher WM capacity were better at suppressing task-irrelevant auditory distractors (Conway, Cowan, & Bunting, 2001; Nagaraj, Magimairaj, & Schwartz, 2020). In ERPs, the mismatch negativity (MMN) is elicited by infrequent (deviant) sounds within repetitive (standard) sounds, and its amplitude reflects neural sensitivity to auditory change (Risto Näätänen, 1990). Our recent work (Kong et al., in press) demonstrated that when task-irrelevant sounds were presented simultaneously visual stimuli, children with ADHD showed greater MMN amplitudes than TD children, contrasting with the attenuated MMN responses in ADHD reported by Yang et al. (2024) using task-relevant auditory distractors. Yang et al. (2024) further reported that stronger neural responses to task-relevant auditory distractors correlated with higher visual WM capacity in TD children but with lower capacity in ADHD, suggesting that the relationship between auditory distraction and visual WM capacity may depend on both distractor relevance and individual differences in attentional control.

Despite these findings, it remains unclear whether and how susceptibility to task-irrelevant auditory distractors is related to visual WM capacity in children with and without ADHD. Individuals with different WM capacities differ in how they allocate attention (Zhong et al., 2024), and such allocation across sensory modalities can be influenced by audiovisual competition (Huang, Wang, & Zhang, 2024). Therefore, the level of audiovisual competition may shape the relationship between visual WM capacity and neural susceptibility to task-irrelevant auditory distraction. We thus examined the potential links between visual WM capacity and neural susceptibility to auditory distraction in children with and without ADHD and tested whether they are modulated by the level of audiovisual competition.

In this study, electroencephalography (EEG) signals were recorded while the children performed visual tasks with simultaneous task-irrelevant auditory distractors across two experiments with different levels of audiovisual competition. We used MMN amplitude as a neural index of task-irrelevant auditory distractor susceptibility and K-score as a behavioral measure of visual WM capacity. Given that TD children can flexibly modulate neural responses to task-irrelevant distraction depending on attentional demands (Kong et al., in press), we expected that the audiovisual competition level would influence the relationship between visual WM capacity and auditory distractor susceptibility. For TD children, we considered two possibilities: those with higher visual WM capacity might show either weaker neural responses to distractors (smaller MMN amplitudes) because of more efficient focusing, or stronger responses (larger MMN amplitudes) due to broader attention across modalities, depending on competition level. In contrast, children with ADHD may have limited cognitive resources that hinder the filtering of irrelevant input and reduce attentional flexibility (Barkley, 1997; Sonuga-Barke, Wiersema, van der Meere, & Roeyers, 2010), leading us to hypothesize that those with lower WM capacity would always show greater susceptibility to auditory distraction, with this relationship less sensitive to audiovisual competition.

## Methods

### Participants

A total of 186 children were initially screened for eligibility, and 160 children (Experiment 1: 45 ADHD and 53 TD; Experiment 2: 27 ADHD and 35 TD) participated in this study. All children with ADHD were recruited from Peking University Sixth Hospital/Institute of Mental Health, and TD children were enrolled from local primary schools. ADHD diagnoses were established through semistructured interviews conducted by qualified psychiatrists using the Kiddie Schedule for Affective Disorders and Schizophrenia for School-Age Children-Present and Lifetime Version (K-SADS-PL; Kaufman et al., 1997) according to the DSM-V criteria. The symptom severity was rated by parents on the ADHD Rating Scale (ADHD-RS-IV; DuPaul, Power, Anastopoulos, & Reid, 1998).

All children met the following inclusion criteria: (a) normal hearing and normal or corrected-to-normal vision, (b) no history of neurological disorders or other severe diseases, and (c) full-scale IQ above 80 assessed by the Wechsler Intelligence Scale for Children (WISC-IV; Wechsler, 2003). Based on these criteria, 26 children (Experiment 1: 12 ADHD and 3 TD; Experiment 2: 9 ADHD and 2 TD) were excluded because of comorbidities or psychiatric concerns. Data collection was terminated early for two children with ADHD and one TD child at their own request in Experiment 1. Data from an additional 11 children (Experiment 1: 3 ADHD and 2 TD; Experiment 2: 4 ADHD and 2 TD) were excluded because of excessive artifacts. A recruitment and exclusion flowchart is presented in Figure 1. The final sample included 90 children from Experiment 1 (ADHD: 8 girls and 32 boys, age = 10.57 ± 2.16; TD: 15 girls and 35 boys, age = 9.86 ± 1.49) and 56 children from Experiment 2 (ADHD: 4 girls and 19 boys, age = 10.47 ± 1.86; TD: 7 girls and 26 boys, age = 10.11 ± 1.62).

**Figure 1.**
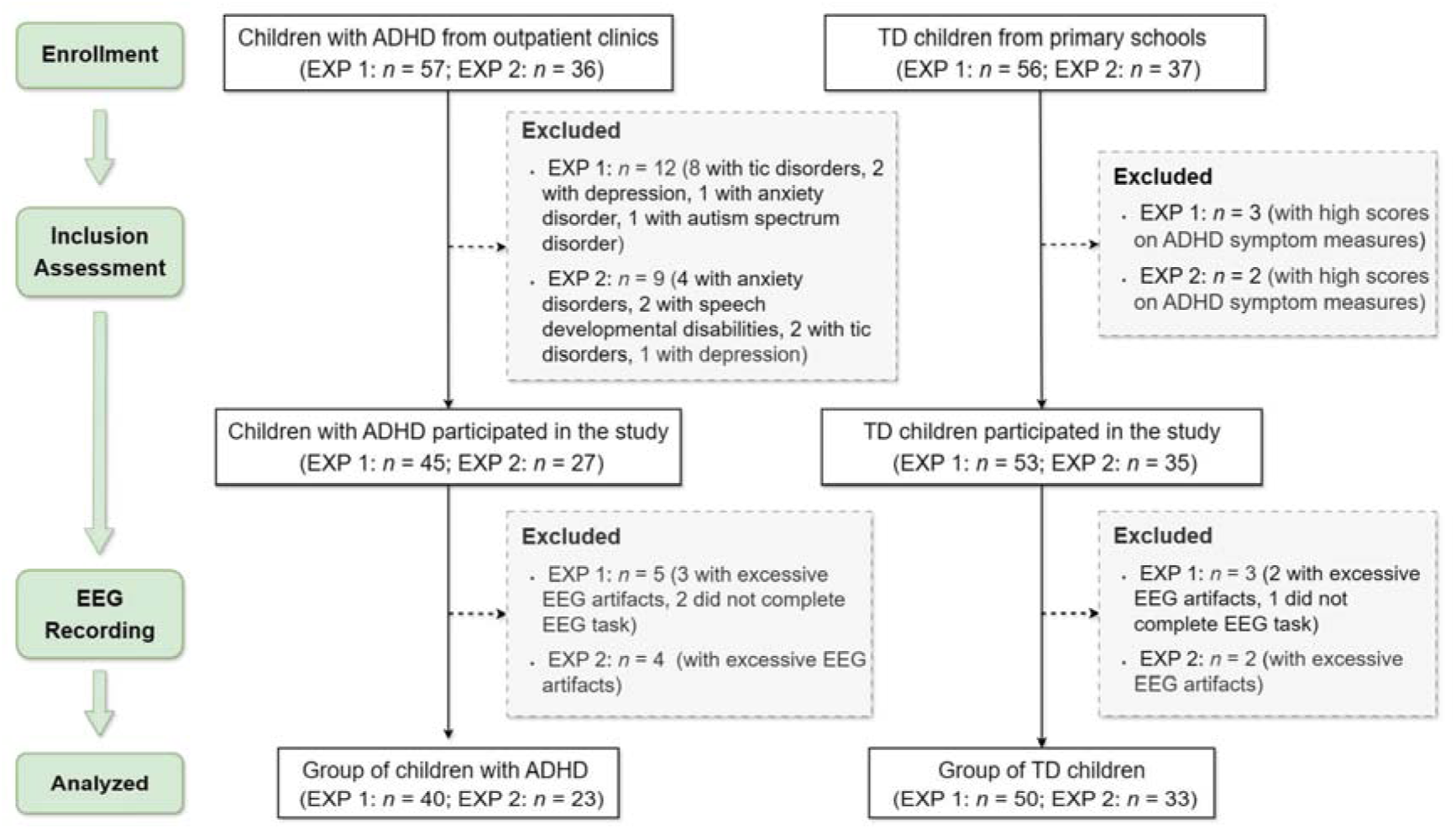
Flowchart of subject enrollment. ADHD, attention-deficit/hyperactivity disorder; TD, typically developing.

A post hoc power analysis using G*Power (Faul, Erdfelder, Lang, & Buchner, 2007) indicated that our sample sizes achieved a sufficient power above 0.45 to detect a medium effect size (*f* = 0.25, α = .05). Moreover, the sample sizes were larger than those reported in some MMN studies of children (Bruder et al., 2011; Yang et al., 2022). No significant group differences were observed in age, sex, handedness, or IQ in either experiment (*p*s > .084; Table 1). This study was approved by the local Ethics Committee, and informed consent was obtained from all participants according to the Declaration of Helsinki.

**Table 1.**
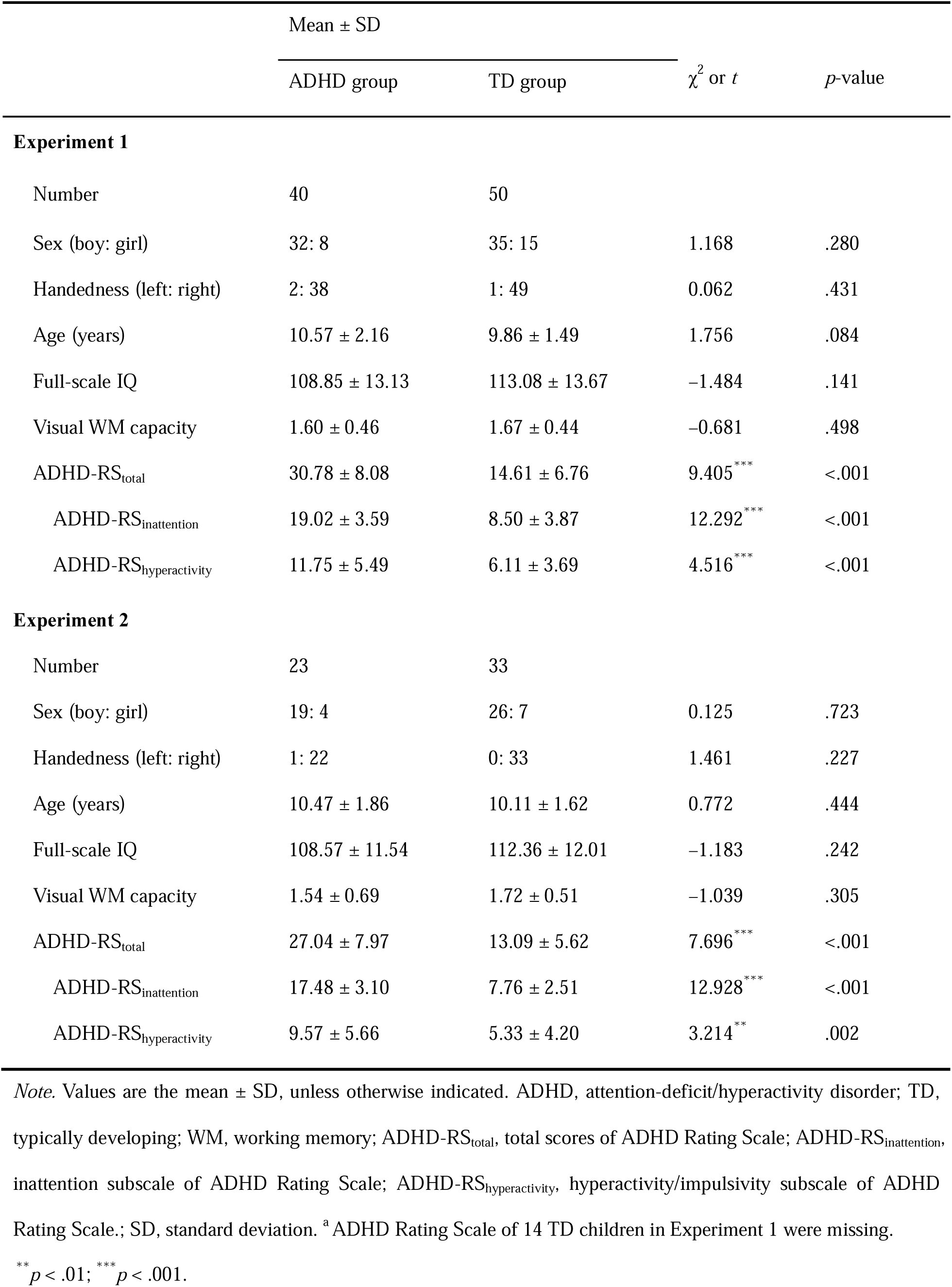
Demographic information of the ADHD and TD groups in the final sample.

| | Mean ± SD | | $\chi^2$ or $t$ | $p$-value |
| --- | --- | --- | --- | --- |
|  | ADHD group | TD group |  |  |
| Experiment 1 |  |  |  |  |
| Number | 40 | 50 |  |  |
| Sex (boy: girl) | 32: 8 | 35: 15 | 1.168 | .280 |
| Handedness (left: right) | 2: 38 | 1: 49 | 0.062 | .431 |
| Age (years) | 10.57 ± 2.16 | 9.86 ± 1.49 | 1.756 | .084 |
| Full-scale IQ | 108.85 ± 13.13 | 113.08 ± 13.67 | −1.484 | .141 |
| Visual WM capacity | 1.60 ± 0.46 | 1.67 ± 0.44 | −0.681 | .498 |
| ADHD-RS<sub>total</sub> | 30.78 ± 8.08 | 14.61 ± 6.76 | 9.405<sup>***</sup> | <.001 |
| ADHD-RS<sub>inattention</sub> | 19.02 ± 3.59 | 8.50 ± 3.87 | 12.292<sup>***</sup> | <.001 |
| ADHD-RS<sub>hyperactivity</sub> | 11.75 ± 5.49 | 6.11 ± 3.69 | 4.516<sup>***</sup> | <.001 |
| Experiment 2 |  |  |  |  |
| Number | 23 | 33 |  |  |
| Sex (boy: girl) | 19: 4 | 26: 7 | 0.125 | .723 |
| Handedness (left: right) | 1: 22 | 0: 33 | 1.461 | .227 |
| Age (years) | 10.47 ± 1.86 | 10.11 ± 1.62 | 0.772 | .444 |
| Full-scale IQ | 108.57 ± 11.54 | 112.36 ± 12.01 | −1.183 | .242 |
| Visual WM capacity | 1.54 ± 0.69 | 1.72 ± 0.51 | −1.039 | .305 |
| ADHD-RS<sub>total</sub> | 27.04 ± 7.97 | 13.09 ± 5.62 | 7.696<sup>***</sup> | <.001 |
| ADHD-RS<sub>inattention</sub> | 17.48 ± 3.10 | 7.76 ± 2.51 | 12.928<sup>***</sup> | <.001 |
| ADHD-RS<sub>hyperactivity</sub> | 9.57 ± 5.66 | 5.33 ± 4.20 | 3.214<sup>**</sup> | .002 |

*Note.* Values are the mean ± SD, unless otherwise indicated. ADHD, attention-deficit/hyperactivity disorder; TD, typically developing; WM, working memory; ADHD-RS<sub>total</sub>, total scores of ADHD Rating Scale; ADHD-RS<sub>inattention</sub>, inattention subscale of ADHD Rating Scale; ADHD-RS<sub>hyperactivity</sub>, hyperactivity/impulsivity subscale of ADHD Rating Scale.; SD, standard deviation. <sup>a</sup> ADHD Rating Scale of 14 TD children in Experiment 1 were missing.
<sup>\*\*</sup> $p < .01$ ; <sup>\*\*\*</sup> $p < .001$ .

### Stimuli and procedure

All experiments were performed in a dimly lit, sound-attenuated and electrically shielded room. Participants were seated alone with their eyes level and directly looking at the fixation cross from a monitor positioned ∼65 cm away. They first performed a behavioral change detection task measuring visual WM capacity and then performed one of the two EEG experiments.

#### Visual WM measures

The change detection task was similar to that in previous WM research (Yang et al., 2024). The task consisted of two sessions, with set sizes of two (*N* = 2) and four (*N* = 4). Each trial began with a memory array containing either two or four colored squares (1.16° × 1.16°) randomly positioned in an 8° × 8° area (Figure S1). The colors of all squares are listed in Figure S2. The memory array was presented for 200 ms, followed by a 900-ms interval. Then, a test probe (single-colored square) appeared at the same location as one of the squares in the memory array. Participants were instructed to take their time in responding after the test probe appeared and to decide whether the color of the test probe match the same-location square in the memory array. Each session included 40 trials, preceded by 10 practice trials to ensure that the participants fully understood the task instructions. The K-score was calculated using the standard formula:

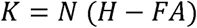

*N*, *H* and *FA* represent the set size, hit rate, and false alarm rate, respectively (Zhong et al., 2024). For each subject, the average K-score of both set sizes was defined as individual visual WM capacity.

#### EEG experiments

In Experiment 1 (Figure 2a), a fixation cross was displayed on a black background. In a random 90% of the trials, the cross remained gray but turned red in the remaining trials (10%). The color never changed in two consecutive trials. Simultaneously, auditory oddball stimuli (10 ms rise and fall times) were presented binaurally through headphones, each lasting 200 ms and followed by a 600-ms interval. The auditory series comprised 80% standard tones and 20% deviant tones, with frequencies counterbalanced across participants between 200 and 800 Hz. The order of tones was randomized, with the restrictions that each deviant tone was preceded by at least one standard tone. The volume of auditory stimuli was individually adjusted to a comfortable level (∼70 dB SPL). Participants were told to ignore the tones and respond as quickly and accurately as possible when the cross turned red. Experiment 1 contained ten 100-trial blocks and lasted ∼ 30 minutes.

**Figure 2.**
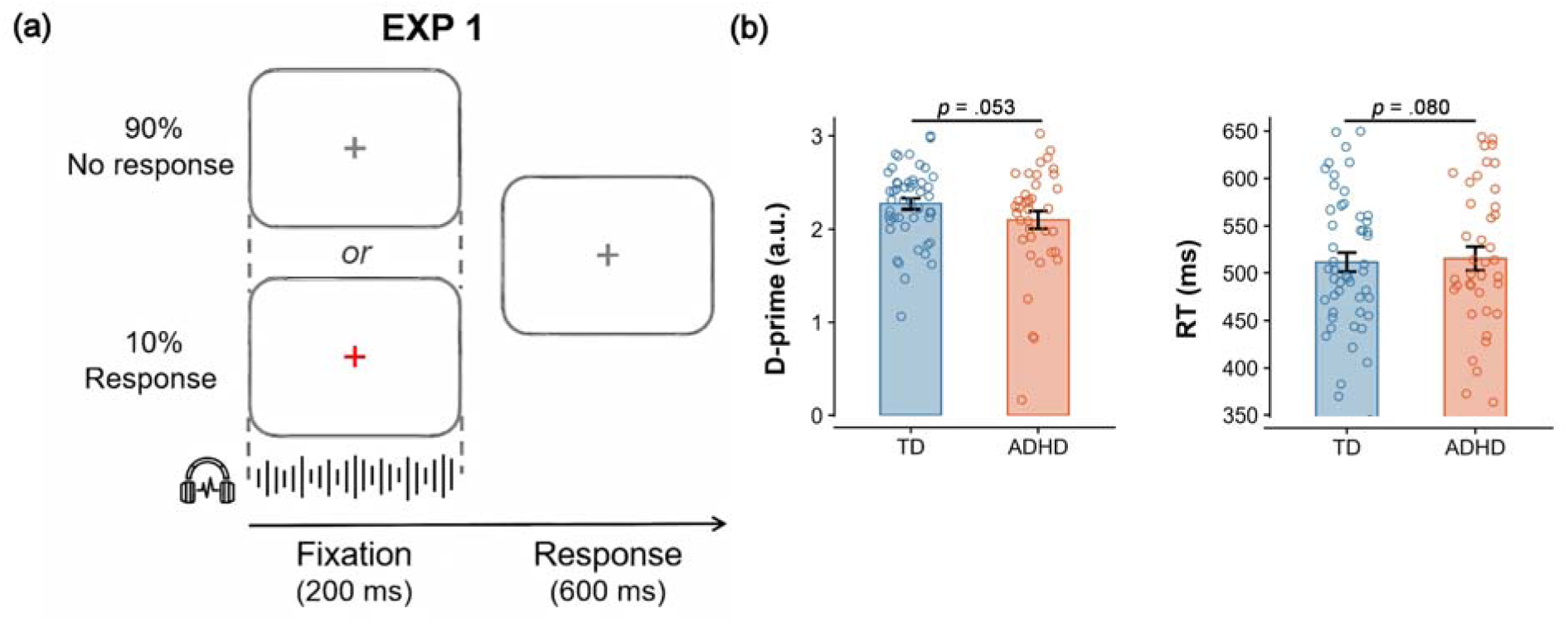
Task paradigm and behavior performance for Experiment 1. (a) The fixation cross remained gray in 90% of the trials, whereas it randomly turned red in the remaining 10% of the trials. Simultaneously, each auditory tone was presented for 200 ms at 600-ms intervals. Participants were instructed to report when the cross turned red (10% of trials, randomly distributed) while ignoring the task-irrelevant auditory stimuli. (b) The d-prime and RT for TD and ADHD groups. The dots denote each subject, and the black error bars represent the standard error of the mean (SEM). D-prime, the aggregate of the proportion of correct hits to false alarms; a larger d-prime value indicates greater sensitivity in detecting whether the gray cross turns red. RT, reaction time.

In Experiment 2 (Figure 4a), we used the same high-demand visual target detection paradigm as in our previous study (Kong et al., in press), where concurrent sounds proved more distracting under a demanding visual condition. 12 items (3.4° × 3.4°) generated a circular search array at a visual angle of 9.2° from the cross on a black screen. These items contained one circle and eleven diamonds. The circle appeared randomly at 2, 4, 8, or 10 o’clock with equal probability. In 90% of the trials, all the items remained light gray, and no response was needed. In the remaining 10% of the trials, the circle turned dark gray, differing from the light gray diamonds. The color never changed in two consecutive trials. Each trial started with a 200-ms presentation of the search array followed by a 600-ms interval. The concurrent auditory tones were identical to those in Experiment 1 and delivered binaurally via headphones. Participants were instructed to ignore task-irrelevant tones and press the “1” key as quickly and accurately as possible when the circle turned dark gray. Experiment 2 lasted ∼30 minutes and comprised eight 100-trials blocks.

### EEG recordings and analysis

EEG signals were collected using a 128-channel EGI system (Electrical Geodesics, Inc., Eugene, OR, USA), with Cz as the online reference and electrode impedance kept below 50 KΩ. EEG data were amplified with a 0.01–400 Hz bandpass and digitized at 1000 Hz. Offline EEG analyses were conducted in EEGLAB (Delorme & Makeig, 2004) using the same methods for both experiments.

To improve data quality, 20 outermost electrodes were excluded because of their susceptibility to motion artifacts (Meyer, Brezack, & Woodward, 2024). EEG data were then downsampled to 250 Hz, filtered at 0.5–30 Hz, and algebraically rereferenced to the average of left and right mastoids. After interpolation of bad electrodes and manual rejection of artifacts, EEG data were epoched from –200 to 600 ms time-locked to the stimulus onset, with the 100-ms prestimulus period serving as baseline. Independent component analysis was applied to reject components with eye movements. Epochs exceeding ± 100 µV in any channel were removed. To mitigate the influence of visual alterations and key presses, trials requiring a response and their subsequent trials were discarded.

The auditory MMN was isolated using difference waves derived by subtracting the ERPs to standard tones from those to deviant tones. The MMN was measured from 10 fronto-central channels surrounding FCz (E4, E5, E6, E11, E12, E13, E19, E20, E112, and E118), which are similar to those used in previous MMN studies (Cantonas et al., 2021; Liang et al., 2025). The average responses within 100–250 ms were calculated to verify the significant presence of the MMN (R. Näätänen, Paavilainen, Rinne, & Alho, 2007). To account for individual latency variations. the MMN amplitude for each participant was calculated as the mean value of a 40-ms window centered at the individual peak (the most negative value in a 100-ms window around the group-averaged peak time). The MMN latency was the time at which the difference waveform reached 50% of its individual peak amplitude within a 140-ms window centered at the group-averaged peak time.

### Statistical analysis

Statistical analyses were performed using SPSS (IBM Inc., Armonk, NY, USA). Chi-square tests and independent-sample *t* tests were used to compare group differences in discontinuous (handedness, sex) and continuous (age, IQ, ADHD symptom scores) demographical variables, respectively. One-sample *t* tests were applied to evaluate whether the MMN significantly differed from zero. One-way ANCOVAs with age as a covariate were conducted to assess whether the two groups differed in behavioral indicators (d-prime, reaction time [RT]) and electrophysiological indices (MMN amplitude, MMN latency). Partial correlation analyses were used to investigate the associations between MMN amplitude and K-score, with age as a covariate to rule out the influence of age on ERPs (Böttger, Herrmann, & von Cramon, 2002). As exploratory analyses, we examined the relationship between MMN amplitude and ADHD symptom scores using partial correlations with age as a covariate. Moreover, to compare group differences in auditory MMN amplitude and K-score between the two experiments, we further performed two-way ANCOVAs using age as a covariate with experiment (Experiment 1, Experiment 2) and group (ADHD, TD) as factors. The exploratory results can be seen in Appendix S3, S4 and S5. All significance levels were set at *p* < .05.

## Results

### Experiment 1

In Experiment 1, the audiovisual competition was set to a low level. We investigated whether susceptibility to auditory distractors was associated with visual WM capacity in children with and without ADHD. The participants first completed a behavioral visual change detection task to measure K-score. EEG signals were recorded when the participants performed a visual target detection task that required monitoring a fixation cross while ignoring concurrent auditory tones, which elicited the MMN. We then examined the correlations between MMN amplitude and K-score in each group.

#### Behavioral Results

In the behavioral change detection task, one-way ANCOVA with age as a covariate was conducted on the K-scores to compare visual WM capacity between the two groups. No significant group difference in visual WM capacity was found (*F*_(1,87)_ = 1.948, *p* = .166, *η*_*p*_^2^ = 0.022; Table 1).

In the visual target detection task during EEG recording, we analyzed d-prime (a sensitivity index from signal detection theory; Hautus, Macmillan, & Creelman, 2021) and RT for correct trials to examine group differences in behavioral performance. The results of one-way ANCOVAs (Figure 2b) revealed borderline significant group differences in d-prime (*F*_(1,87)_ = 3.854, *p* = .053, *η*_*p*_^2^ = 0.042) and RT (*F*_(1,87)_ = 3.128, *p* = .080, *η*_*p*_^2^ = 0.035).

#### Auditory MMN

A negative difference between the standard and deviant auditory tones was observed at the fronto-central electrode in both groups of children (Figure 3a). One-sample *t* tests confirmed that the auditory MMN amplitudes were significantly different from zero in both groups (ADHD: –2.10 ± 1.78 μV, *t*_(39)_ = –7.467, *p* < .001, Cohen’s *d* = –1.181; TD: –2.17 ± 1.74 μV, *t*_(49)_ = –8.843, *p* < .001, Cohen’s *d* = –1.251), indicating reliable neural responses to the auditory stimuli despite instructions to ignore them. Further one-way ANCOVAs (Figure 3b) revealed no significant group difference in auditory MMN amplitudes (*F*_(1,87)_ = 0.503, *p* = .480, *η*_*p*_^2^ = 0.006) or latencies (*F*_(1,87)_ = 1.602, *p* = .209, *η*_*p*_^2^ = 0.018), suggesting comparable sensitivity for auditory distractors in children with and without ADHD (M.-T. Yang et al., 2015). Topographic maps revealed the classic fronto-central distribution of the MMN component for both groups (Figure 3c).

**Figure 3.**
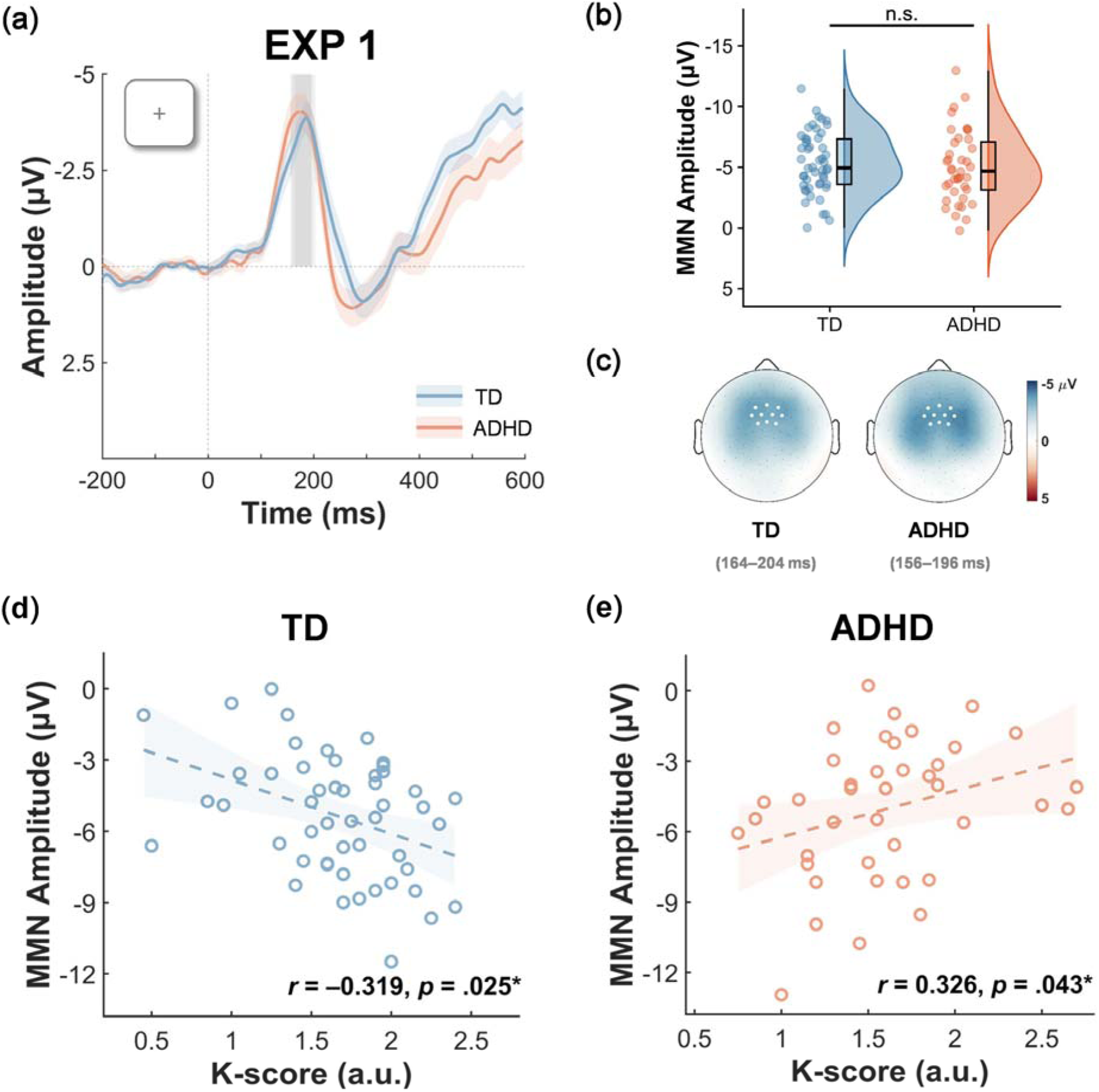
MMN and correlation results for Experiment 1. (a) Grand average of deviant-minus-standard difference waves for the TD (blue) and ADHD (orange) groups. The vertical dotted line at 0 ms indicates the time of stimulus onset. The shading in the wave plots represents the standard error of each group. The gray rectangular areas represent the time windows used to generate MMN topographical maps for the two groups. (b) The MMN amplitudes for the two groups. Raincloud plots show the data distribution (density), box plots, and individual data points. Box plots indicate the interquartile ranges and medians. (c) The MMN scalp topographies for the two groups. The white dots refer to the electrodes from which the ERP waveforms were derived. (d) Correlation between K-score and MMN amplitude in TD children and (e) children with ADHD. The shading in the correlation plots represents the 95% confidence intervals (CIs) of the trend line. n.s., not significant. \**p*□<□.05.

#### Correlation between auditory MMN and visual WM capacity

To further examine the relationship between susceptibility to task-irrelevant auditory distractors and visual WM capacity, we conducted partial correlation analyses between the auditory MMN amplitude and K-score, controlling for age. The results revealed opposite association patterns between MMN amplitude and K-score across groups. In TD children, higher K-scores were significantly associated with smaller MMN amplitudes (*r* = –0.319, *p* = .025; Figure 3d), suggesting that children with better visual WM capacity showed weaker neural responses to task-irrelevant auditory distractors. In contrast, in children with ADHD, higher K-scores were significantly correlated with larger MMN amplitudes (*r* = 0.326, *p* = .043; Figure 3e), indicating that better visual WM capacity was linked with greater susceptibility to task-irrelevant sounds. These contrasting patterns across groups are consistent with previous findings in Yang et al. (2024), although we used task-irrelevant rather than task-relevant auditory distractors. These results suggest a close relationship between visual WM capacity and distractor susceptibility, which differs in ADHD and TD children.

### Experiment 2

In Experiment 2, we further increased the visual demand to make the auditory distractors more disruptive, thereby exploring whether the previously observed associations in Experiment 1 would be altered. The auditory stimuli were identical across two experiments, which enabled a direct comparison of MMN responses between Experiments 1 and 2. EEGs were recorded when participants performed a new visual target detection task with concurrent auditory distractors, where the audiovisual competition increased. As in Experiment 1, we measured the visual WM capacity and examined the relationships between auditory MMN amplitude and K-score.

#### Behavioral Results

In the behavioral change detection task, the results (Table 1) revealed a borderline significant group difference in visual WM capacity (*F*_(1,53)_ = 3.417, *p* = .070, *η*_*p*_^2^ = 0.061). Although neither experiment showed significant group differences separately, further cross-experiment comparison revealed that the ADHD group exhibited significantly lower visual WM capacity than the TD group (see Appendix S5).

In the visual target detection task during EEG recording, children with ADHD demonstrated significantly lower d-prime (*F*_(1,53)_ = 18.662, *p* < .001, 1^2^ = 0.260) and longer RT (*F*_(1,53)_ = 32.647, *p* < .001, 1^2^ = 0.381) than TD children did (Figure 4b).

**Figure 4.**
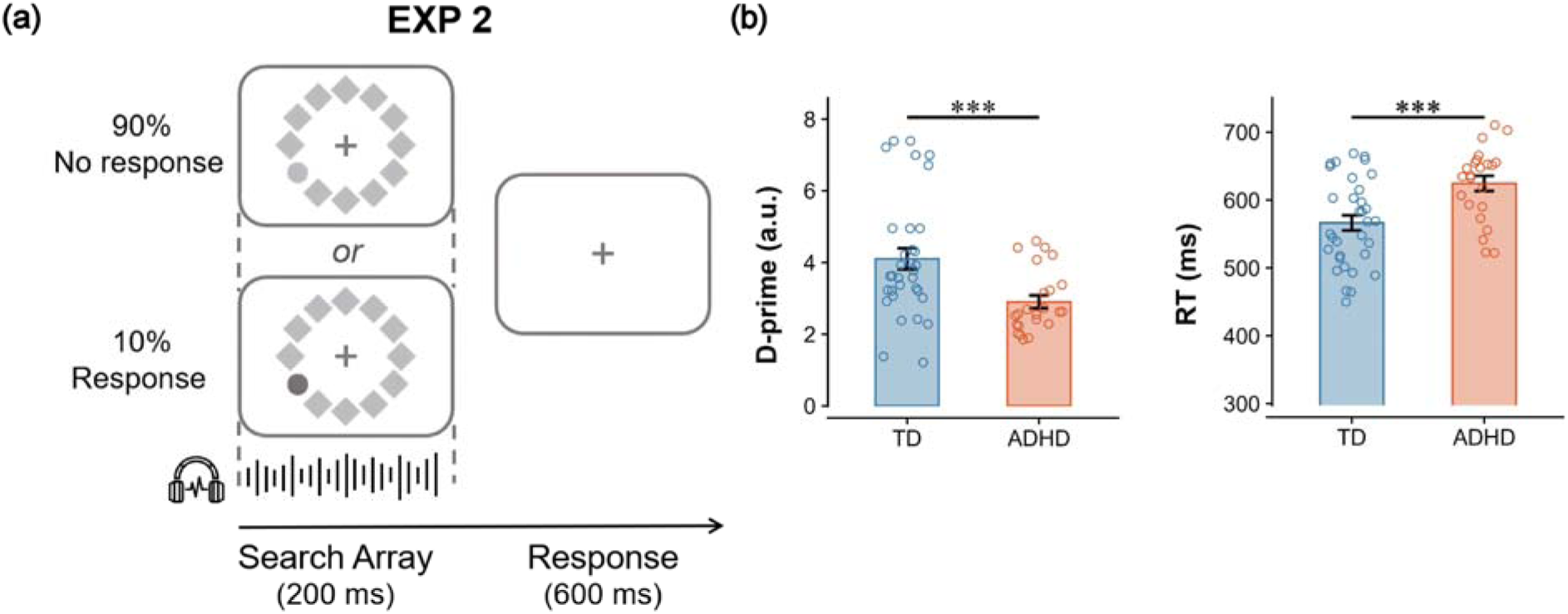
Task paradigm and behavior results for Experiment 2. (a) The visual search array consisted of one gray circle and eleven gray diamonds. The circle remained light gray in 90% of the trials, whereas it randomly turned dark gray in the remaining 10% of the trials. Moreover, the same auditory stimuli as in Experiment 1 were presented in sync with the visual search array. Participants were instructed to report the presence of a dark gray circle while ignoring task-irrelevant auditory tones. (b) The d-prime and RT for TD and ADHD groups. D-prime, aggregate of the proportion of correct hits to false alarms; a larger d-prime value denotes greater sensitivity in detecting whether the light gray circle turns dark gray. RT, reaction time. \*\**p*□<□.01, \*\*\**p*□<□.001.

#### Auditory MMN

The deviant-minus-standard difference was characterized by a negative deflection over the fronto-central areas (Figure 5a). The auditory MMN amplitudes significantly differed from zero in both groups (ADHD: –1.77 ± 2.08 μV, *t*_(22)_ = –4.081, *p* < .001, Cohen’s *d* = –0.851; TD: –0.93 ± 1.52 μV, *t*_(32)_ = –3.527, *p* = .001, Cohen’s *d* = –0.614), confirming a robust MMN response to task-irrelevant auditory distractors. The results (Figure 5b) revealed that, compared with TD children, children with ADHD exhibited significantly greater MMN amplitudes (*F*_(1,53)_ = 4.867, *p* = .032, *η*_*p*_^2^ = 0.084) and shorter latencies (*F*_(1,53)_ = 16.736, *p* < .001, *η*_*p*_^2^ = 0.240), indicating their increased sensitivity to auditory distractors during the synchronous high-demand visual task, which aligns with our prior findings (Kong et al., in press). The MMN topographies for both groups are shown in Figure 5c. Further cross-experiment comparison revealed smaller MMN amplitudes in Experiment 2 than in Experiment 1 for TD children, whereas no significant difference in the MMN amplitudes between the two experiments was found for children with ADHD (see Appendix S5).

**Figure 5.**
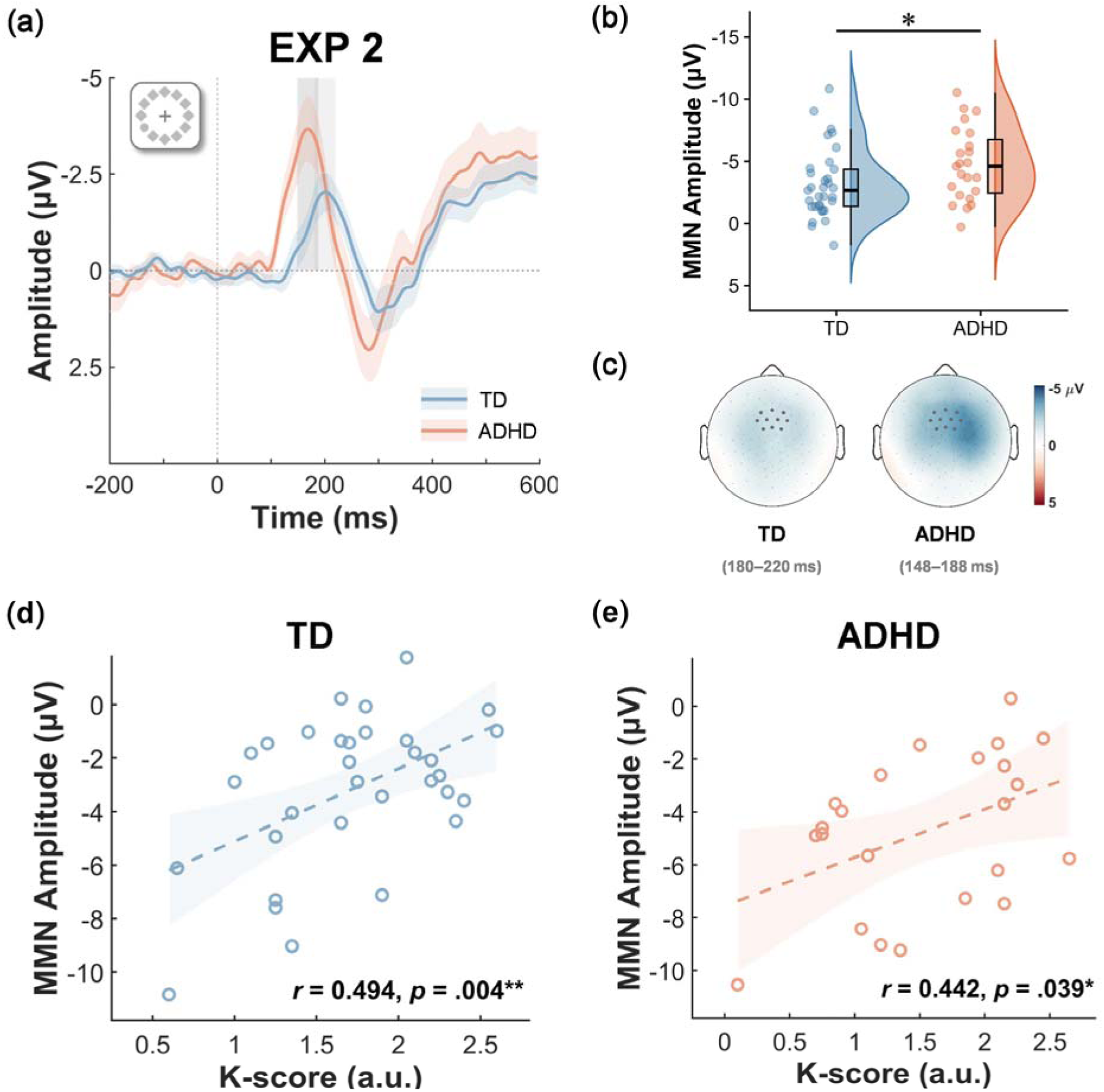
MMN and correlation results for Experiment 2. (a) Grand average of deviant-minus-standard difference waves for the TD (blue) and ADHD (orange) groups. The vertical dotted line at 0 ms indicates the time of stimulus onset. The shading in the wave plots refers to the standard error of each group. The gray rectangular areas represent the time windows used to generate MMN topographical maps for the two groups. (b) The MMN amplitudes for the two groups. Raincloud plots show the data distribution (density), box plots and individual data points. Box plots depict the interquartile ranges and medians. (c) The MMN scalp topographies for the two groups. The black dots refer to the electrodes from which the ERP waveforms were obtained. (d) Correlation between K-score and MMN amplitudes in the TD group and (e) ADHD group. Shading in the correlation plots represents the 95% CIs of the trend line. \**p*□<□.05, \*\**p*□<□.01.

#### Correlation between auditory MMN and visual WM capacity

Correlation results (Figure 5e) revealed that, similar to those in Experiment 1, lower K-scores were significantly associated with greater MMN amplitudes in the ADHD group (*r* = 0.442, *p* = .039). In other words, children with ADHD who had lower visual WM capacity tended to exhibit stronger neural responses to task-irrelevant auditory distractors, regardless of the audiovisual competition level. Similarly, TD children also showed a negative association between MMN amplitude and K-score (*r* = 0.494, *p* = .004; Figure 5d). This pattern was reversed compared with that observed in TD children in Experiment 1. That is, when the audiovisual competition increased, TD children with lower visual WM capacity became more susceptible to auditory distraction. Our findings revealed that under high audiovisual competition, both ADHD and TD groups showed a negative association between visual WM capacity and susceptibility to task-irrelevant auditory distractors.

## Discussion

The present study examined how individual differences in visual WM capacity (as indexed by the K-score) are related to the susceptibility to task-irrelevant auditory distractors (as indexed by the MMN amplitude) in children with and without ADHD. Across two experiments, TD children showed task-dependent patterns: higher visual WM capacity was associated with greater MMN amplitudes under low audiovisual competition in Experiment 1, but this relation reversed under high audiovisual competition in Experiment 2, with lower visual WM capacity predicting greater MMN responses. In contrast, children with ADHD demonstrated a consistent association across experiments: those with lower visual WM capacity were more susceptible to auditory distractors regardless of competition level. These findings highlight the close relationship between children’s susceptibility to task-irrelevant auditory distraction and visual WM capacity, which is jointly influenced by audiovisual competition and attentional control ability and differs between children with and without ADHD.

Group comparisons revealed that in Experiment 1, MMN did not differ between TD and ADHD children, consistent with previous findings (Winsberg, Javitt, & Shanahan/Silipo, 1997; M.-T. Yang et al., 2015), suggesting that group differences in auditory distraction are less pronounced under low visual attentional demand (Kong et al., in press). In contrast, under high audiovisual competition in Experiment 2, children with ADHD exhibited significantly greater MMN responses than TD children. Across experiments, TD children showed flexible responses, whereas children with ADHD consistently overreacted to auditory distractors (Kong et al., in press), suggesting their impairment in filtering task-irrelevant distraction (Schneidt, Jusyte, Rauss, & Schönenberg, 2018). Compared with TD children, children with ADHD showed comparable visual WM capacity in separate experiments, but combining data across experiments revealed lower visual WM capacity, consistent with previous reports of reduced capacity in ADHD (Yang et al., 2024). This suggests that group differences in visual WM may be modest, context-dependent, and obscured by individual variability in smaller samples.

In TD children, the association between visual WM capacity and neural sensitivity to task-irrelevant auditory change depended on the level of audiovisual competition. In Experiment 1 (low audiovisual competition), children with higher visual WM capacity predicted greater auditory MMN amplitudes. This pattern differed from previous visual domain findings showing that adults with higher visual WM capacity better resist task-irrelevant visual distraction (Gaspar et al., 2016). Conversely, it aligns with the positive correlation between visual WM capacity and susceptibility to task-relevant auditory distractors in TD children reported by Yang et al. (2024), who proposed that this reflects a shared neurophysiological mechanism between visual WM and auditory processing when attentional resources are plentiful. Zhong et al. (2024) suggested that the attentional control setting may globally enhance all the stimuli with goal-related features such that adults with high visual WM capacity show stronger capture by task-relevant distractors due to better top-down selection ability. In this study, we found a similar pattern in TD children when the distractors were task-irrelevant rather than relevant. On the basis of load theory (Lavie, 2005), we speculated that this global-enhancement mechanism may also depend on task demand: under low audiovisual competition, the visual task imposes a low demand, leaving attentional resources available to process task-irrelevant distractors. Hence, TD children with higher visual WM capacity may enhance auditory processing more strongly, allowing even irrelevant auditory inputs to receive greater encoding and elicit larger MMN responses.

In contrast, under high audiovisual competition in Experiment 2, TD children demonstrated the opposite pattern: lower visual WM capacity predicted larger MMN amplitudes to the identical task-irrelevant distractors used in Experiment 1. This reversal is naturally explained by invoking a filtering mechanism that becomes necessary when the visual task consumes most attentional resources. Under high audiovisual competition, successful performance may depend on early or proactive suppression of irrelevant distractors (Gaspar et al., 2016). TD children with higher visual WM capacity can better engage suppression processes to limit distractor processing, leading to attenuated MMN amplitudes under higher audiovisual competition. Thus, TD children exhibit a flexible, context-dependent pattern: under low audiovisual competition, abundant resources may allow enhanced encoding of multisensory inputs, leading to a positive visual WM–auditory MMN relationship; under high audiovisual competition, they seem to engage in active suppression of auditory processing and prioritize the demanding visual task, thereby reversing the association.

For children with ADHD, lower visual WM capacity was consistently linked to larger MMN amplitudes across experiments, even under low audiovisual competition. The stability suggests reduced context-dependent flexibility, which is consistent with theories characterizing ADHD as involving constrained and less flexible control of attention (Barkley, 1997; Sonuga-Barke et al., 2010). This pattern may reflect limited cognitive resources in children with ADHD (Hwang, Gau, Hsu, & Wu, 2010): even under low competition, those with higher visual WM capacity may lack spare resources for global enhancement of task-irrelevant sounds. Moreover, those with lower visual WM capacity may be less able to deploy effective suppression for task-irrelevant auditory distractors. When audiovisual competition increased in Experiment 2, MMN amplitudes were still large in low-capacity children, suggesting that higher competition further amplified their susceptibility to distraction. Overall, children with ADHD follow a more fixed response pattern, with the visual WM–auditory MMN association remaining negative across different contexts.

We found that the larger MMN amplitude did not significantly explain the greater severity of ADHD symptoms in either experiment, contrasting with studies reporting significant associations between them (Yang et al., 2024). One possible explanation is differences in task design or stimulus characteristics, which may influence the neural processes captured by the MMN. In support of our findings, other work suggests that attentional capture by irrelevant or external distraction may not predict ADHD symptom severity (Coll-Martín, Carretero-Dios, & Lupiáñez, 2021), whereas internal distraction such as mind wandering or intrusive thoughts shows a closer link (Osborne, Zhang, Carlson, Shah, & Jonides, 2023). Individual differences, such as the activation of the default attention network, may also affect the ability to suppress distractors in ADHD (Fassbender et al., 2009). Further research should examine whether these individual differences, along with task and stimulus characteristics, modulate the relationship between susceptibility to task-irrelevant auditory distractors and ADHD symptom severity.

This study contributes to a broader understanding of how the relationship between visual WM capacity and auditory distraction is influenced by audiovisual competition and how these dynamics contribute to our knowledge of children with ADHD. By demonstrating that TD children flexibly shift between global enhancement and proactive suppression mechanisms depending on attention demand, whereas children with ADHD show a rigid and consistently negative association across contexts, our findings highlight the importance of considering both individual differences and task context. These findings support the theoretical accounts of ADHD, suggesting that deficits may involve both less flexible attentional control and executive function impairments, such as difficulties in inhibiting inappropriate responses (Barkley, 1997; Sonuga-Barke et al., 2010).

Nonetheless, several limitations of this study should be noted. The sample size in Experiment 2 was relatively small, and larger samples would help to confirm the observed effects. More generally, MMN amplitude may depend on the diversity of auditory stimuli and task design (R. Näätänen et al., 2007); future studies could vary auditory distractor types and visual task difficulty to increase diversity and generalizability. In addition, the between-subject design limited within-individual comparisons; follow-up research could use a within-subject design to better compare these relationships under different contexts.

## Conclusions

Our study provides neurophysiological evidence that children’s susceptibility to task-irrelevant auditory distractors is closely linked to visual WM capacity, with distinct patterns observed in the TD and ADHD groups. In TD children, this relationship is flexible and context-dependent, whereas in children with ADHD, it remains consistently negative across different levels of audiovisual competition. These findings shed light on the neural mechanisms underlying attentional control and distraction in ADHD patients and highlight the role of WM capacity in modulating the impact of irrelevant auditory input.

## Ethical considerations

Ethical approval was granted by the Institutional Review Boards of Peking University Sixth Hospital/Institute of Mental Health and Beijing Normal University. Both written and verbal informed consent was obtained from all the participants and their caregivers in accordance with the Declaration of Helsinki.

## Abbreviations

ADHD: attention-deficit/hyperactivity disorder
ANCOVA: analysis of covariance
EEG: electroencephalography
ERP: event-related potential
MMN: mismatch negativity
RT: reaction time
SD: standard deviation
TD: typically developing
WM: working memory.

## Acknowledgements

The present research was supported by the STI 2030—Major Projects (No.2021ZD0200500 to Y.S.), the National Natural Science Foundation of China (No. 32271094 to Y.S.; No. 82371549 to L.S.). We thank all the participants and their caregivers for their involvement in our experiments. All authors reported no biomedical financial interests or potential conflicts of interest.

## Data availability statement

Data supporting the findings of this study are available from the corresponding authors upon reasonable request. This study was not preregistered.

## Key points

- Children with attention-deficit/hyperactivity disorder (ADHD) showed increased distractibility; in adults, this distractibility has been linked to visual working memory (WM).
- How neural responses to task-irrelevant auditory distractors relates to visual WM capacity in children with and without ADHD remains unclear.
- We examined the associations between mismatch negativity (MMN) amplitude and K-score, using electroencephalography (EEG) during visual tasks of varying difficulty with auditory distractors.
- ADHD group showed a stable negative relationship that lower visual WM capacity predicted greater susceptibility, whereas typically developing (TD) children displayed opposite patterns under low versus high audiovisual competition.
- These findings highlight that WM-linked neural susceptibility to auditory distraction may help understand attentional control in ADHD and inform future research on interventions.

## References

American Psychiatric Association. (2013). Diagnostic and statistical manual of mental disorders (5th ed.). Arlington: American Psychiatric Publishing, Inc.

Barkley, R. A. (1997). Behavioral inhibition, sustained attention, and executive functions: Constructing a unifying theory of ADHD. Psychological Bulletin, 121, 65–94.

Biederman, J., Petty, C., Fried, R., Fontanella, J., Doyle, A. E., Seidman, L. J., & Faraone, S. V. (2006). Impact of psychometrically defined deficits of executive functioning in adults with attention deficit hyperactivity disorder. American Journal of Psychiatry, 163, 1730–1738.

Böttger, D., Herrmann, C. S., & von Cramon, D. Y. (2002). Amplitude differences of evoked alpha and gamma oscillations in two different age groups. International Journal of Psychophysiology, 45, 245–251.

Bruder, J., Leppänen, P. H. T., Bartling, J., Csépe, V., Démonet, J., & SchultelKörne, G. (2011). Children with dyslexia reveal abnormal native language representations: Evidence from a study of mismatch negativity. Psychophysiology, 48, 1107–1118.

Burgoyne, A. P., & Engle, R. W. (2020). Attention control: A cornerstone of higher-order cognition. Current Directions in Psychological Science, 29, 624–630.

Cantonas, L.-M., Mancini, V., Rihs, T. A., Rochas, V., Schneider, M., Eliez, S., & Michel, C. M. (2021). Abnormal auditory processing and underlying structural changes in 22q11.2 deletion syndrome. Schizophrenia Bulletin, 47, 189–196.

Coll-Martín, T., Carretero-Dios, H., & Lupiáñez, J. (2021). Attentional networks, vigilance, and distraction as a function of attention-deficit/hyperactivity disorder symptoms in an adult community sample. British Journal of Psychology, 112, 1053–1079.

Conway, A. R. A., Cowan, N., & Bunting, M. F. (2001). The cocktail party phenomenon revisited: The importance of working memory capacity. Psychonomic Bulletin & Review, 8, 331–335.

Delorme, A., & Makeig, S. (2004). EEGLAB: An open source toolbox for analysis of single-trial EEG dynamics including independent component analysis. Journal of Neuroscience Methods, 134, 9–21.

DuPaul, G. J., Power, T. J., Anastopoulos, A. D., & Reid, R. (1998). ADHD rating scale—IV: Checklists, norms, and clinical interpretation (1st ed.). New York: New York: The Guilford Press.

Fassbender, C., Zhang, H., Buzy, W. M., Cortes, C. R., Mizuiri, D., Beckett, L., & Schweitzer, J. B. (2009). A lack of default network suppression is linked to increased distractibility in ADHD. Brain Research, 1273, 114–128.

Faul, F., Erdfelder, E., Lang, A.-G., & Buchner, A. (2007). G*power 3: A flexible statistical power analysis program for the social, behavioral, and biomedical sciences. Behavior Research Methods, 39, 175–191.

Gaspar, J. M., Christie, G. J., Prime, D. J., Jolicœur, P., & McDonald, J. J. (2016). Inability to suppress salient distractors predicts low visual working memory capacity. Proceedings of the National Academy of Sciences, 113, 3693–3698.

Hautus, M. J., Macmillan, N. A., & Creelman, C. D. (2021). Detection theory: A user’s guide (3rd ed.). New York: New York: Psychology Press.

Huang, J., Wang, A., & Zhang, M. (2024). The audiovisual competition effect induced by temporal asynchronous encoding weakened the visual dominance in working memory retrieval. Memory, 32, 1069–1082. (world).

Hwang, S., Gau, S. S., Hsu, W., & Wu, Y. (2010). Deficits in interval timing measured by the dualltask paradigm among children and adolescents with attentionldeficit/hyperactivity disorder. Journal of Child Psychology and Psychiatry, 51, 223–232.

Jonkman, L. M., Kenemans, J. L., Kemner, C., Verbaten, M. N., & Van Engeland, H. (2004). Dipole source localization of event-related brain activity indicative of an early visual selective attention deficit in ADHD children. Clinical Neurophysiology, 115, 1537–1549.

Kaufman, J., Birmaher, B., Brent, D., Rao, U., Flynn, C., Moreci, P., … Ryan, N. (1997). Schedule for affective disorders and schizophrenia for school-age children-present and lifetimeversion (K-SADS-PL): Initial reliability and validity data. Journal of the American Academy of Child & Adolescent Psychiatry, 36, 980–988.

Kofler, M. J., Rapport, M. D., Bolden, J., Sarver, D. E., & Raiker, J. S. (2010). ADHD and working memory: The impact of central executive deficits and exceeding storage/rehearsal capacity on observed inattentive behavior. Journal of Abnormal Child Psychology, 38, 149–161.

Kong, Y., Yuan, X., Sun, L., Dang, C., Wang, Y., Huang, J., … Song, Y. (in press). Altered processing of auditory distractions under competing inputs in children with ADHD. Journal of the American Academy of Child & Adolescent Psychiatry.

Lavie, N. (2005). Distracted and confused?: Selective attention under load. Trends in Cognitive Sciences, 9, 75–82.

Liang, M., Peng, P., Liu, J., Wang, Z., Lai, K., Wang, J., … Wang, S. (2025). Tone and vowel perception delay: Long-term effects of late cochlear implant in children with prelingual deafness. Frontiers in Human Neuroscience, 19, 1516931.

Mason, D. J., Humphreys, G. W., & Kent, L. (2005). Insights into the control of attentional set in ADHD using the attentional blink paradigm. Journal of Child Psychology and Psychiatry, 46, 1345–1353.

McNab, F., & Klingberg, T. (2008). Prefrontal cortex and basal ganglia control access to working memory. Nature Neuroscience, 11, 103–107.

Meyer, M., Brezack, N., & Woodward, A. L. (2024). Neural correlates involved in perspective-taking in early childhood. Developmental Cognitive Neuroscience, 66, 101366.

Näätänen, R., Paavilainen, P., Rinne, T., & Alho, K. (2007). The mismatch negativity (MMN) in basic research of central auditory processing: A review. Clinical Neurophysiology, 118, 2544–2590.

Näätänen, Risto. (1990). The role of attention in auditory information processing as revealed by event-related potentials and other brain measures of cognitive function. Behavioral and Brain Sciences, 13, 201–233.

Nagaraj, N. K., Magimairaj, B. M., & Schwartz, S. (2020). Auditory distraction in school-age children relative to individual differences in working memory capacity. Attention, Perception, & Psychophysics, 82, 3581–3593.

Osborne, J. B., Zhang, H., Carlson, M., Shah, P., & Jonides, J. (2023). The association between different sources of distraction and symptoms of attention deficit hyperactivity disorder. Frontiers in Psychiatry, 14, 1173989.

Plebanek, D. J., & Sloutsky, V. M. (2019). Selective attention, filtering, and the development of working memory. Developmental Science, 22, e12727.

Schneidt, A., Jusyte, A., Rauss, K., & Schönenberg, M. (2018). Distraction by salient stimuli in adults with attention-deficit/hyperactivity disorder: Evidence for the role of task difficulty in bottom-up and top-down processing. Cortex, 101, 206–220.

Sonuga-Barke, E. J. S., Wiersema, J. R., van der Meere, J. J., & Roeyers, H. (2010). Context-dependent dynamic processes in attention deficit/hyperactivity disorder: Differentiating common and unique rffects of state regulation deficits and delay aversion. Neuropsychology Review, 20, 86–102.

Thomas, R., Sanders, S., Doust, J., Beller, E., & Glasziou, P. (2015). Prevalence of attention-deficit/hyperactivity disorder: A systematic review and meta-analysis. Pediatrics, 135, e994–e1001.

Wechsler, D. (2003). Wechsler intelligence scale for children (4th ed.). New York: New York: The Psychological Corporation.

Winsberg, B. G., Javitt, D. C., & Shanahan/Silipo, G. (1997). Electrophysiological indices of information processing in methylphenidate responders. Biological Psychiatry, 42, 434–445.

Yang, H., Guo, J., Yin, W., Deng, Y., Fu, T., Huang, S., … Dang, C.-P. (2024). Association between auditory mismatch negativity and visual working memory in school-age children with attention deficit/hyperactivity disorder. Psychological Medicine, 54, 4798–4808.

Yang, M.-T., Hsu, C.-H., Yeh, P.-W., Lee, W.-T., Liang, J.-S., Fu, W.-M., & Lee, C.-Y. (2015). Attention deficits revealed by passive auditory change detection for pure tones and lexical tones in ADHD children. Frontiers in Human Neuroscience, 9, 470.

Yang, Y., Li, Q., Xiao, Y., Liu, Y., Sun, K., Li, B., & Zheng, Q. (2022). Auditory discrimination elicited by nonspeech and speech stimuli in children with congenital hearing loss. Journal of Speech, Language, and Hearing Research, 65, 3981–3995.

Zhong, C., Qu, Z., Yang, N., Sun, M., Wang, Y., & Ding, Y. (2024). Susceptibility to attentional capture by target-matching distractors predicts high visual working memory capacity. Psychological Science, 35, 1203–1216.

