## Appendix S1; Appendix S2; Appendix S3; Appendix S4; Appendix S5 for "Susceptibility to Task-irrelevant Auditory Distractors in Relation to Visual Working Memory in Children With and Without ADHD"

***Supplement Information***

**Appendix S1. The task procedure of visual WM task**

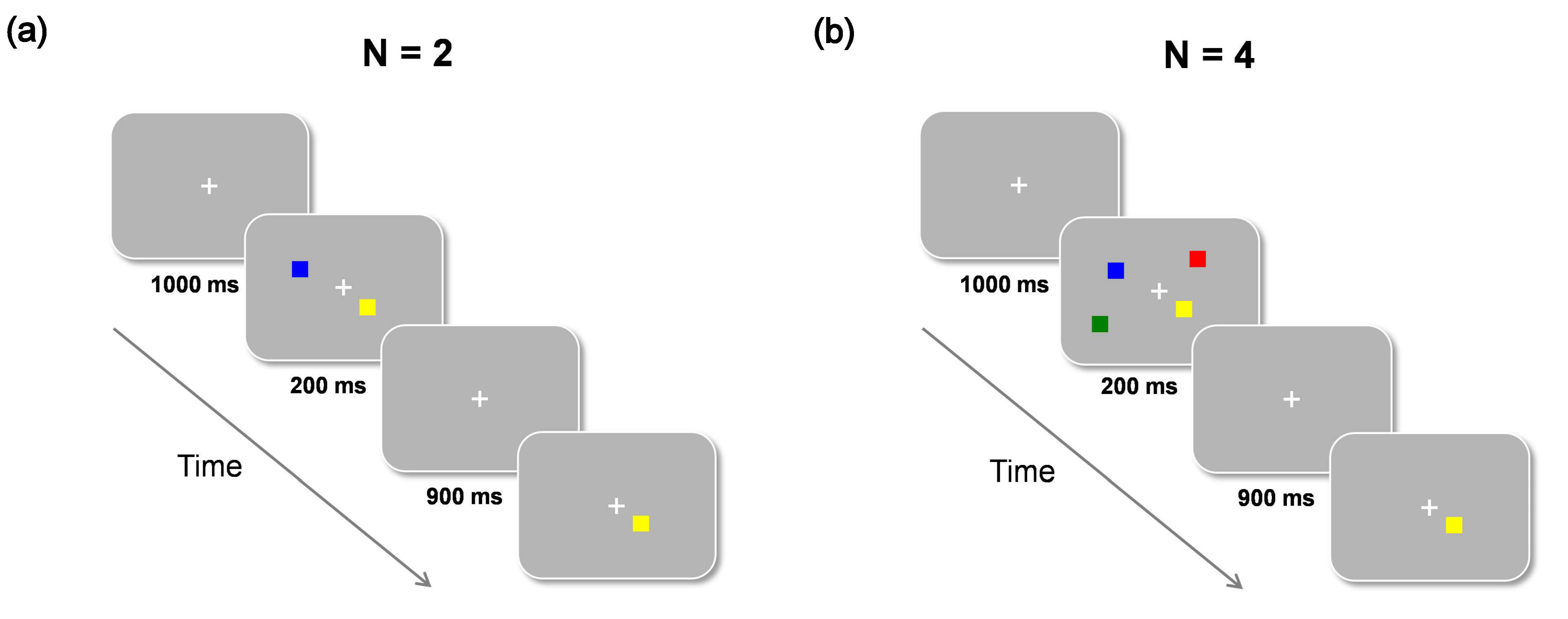

**Figure S1** Schematic diagram of the behavioral change detection task with two set-size sessions: (a) N = 2 and (b) N = 4. Participants are required to determine whether the color of the test probe matches that of the same-location square in the memory array through key presses with “1” for a match or “0” for a nonmatch. N, the number of squares to be remembered.

**Appendix S2. The stimuli colors in visual WM measures**

As illustrated in Figure S2, The colors of all squares were randomly selected without placement from 10 highly distinguishable colors (black [0, 0, 0]; white [255, 255, 255]; red [254, 0, 0]; green [0, 127, 1]; blue [0, 0, 254]; pink [255, 192, 223]; yellow [255, 255, 1]; violet [128, 0, 127]; cyan [1, 255, 255]; orange [255, 102, 0]).

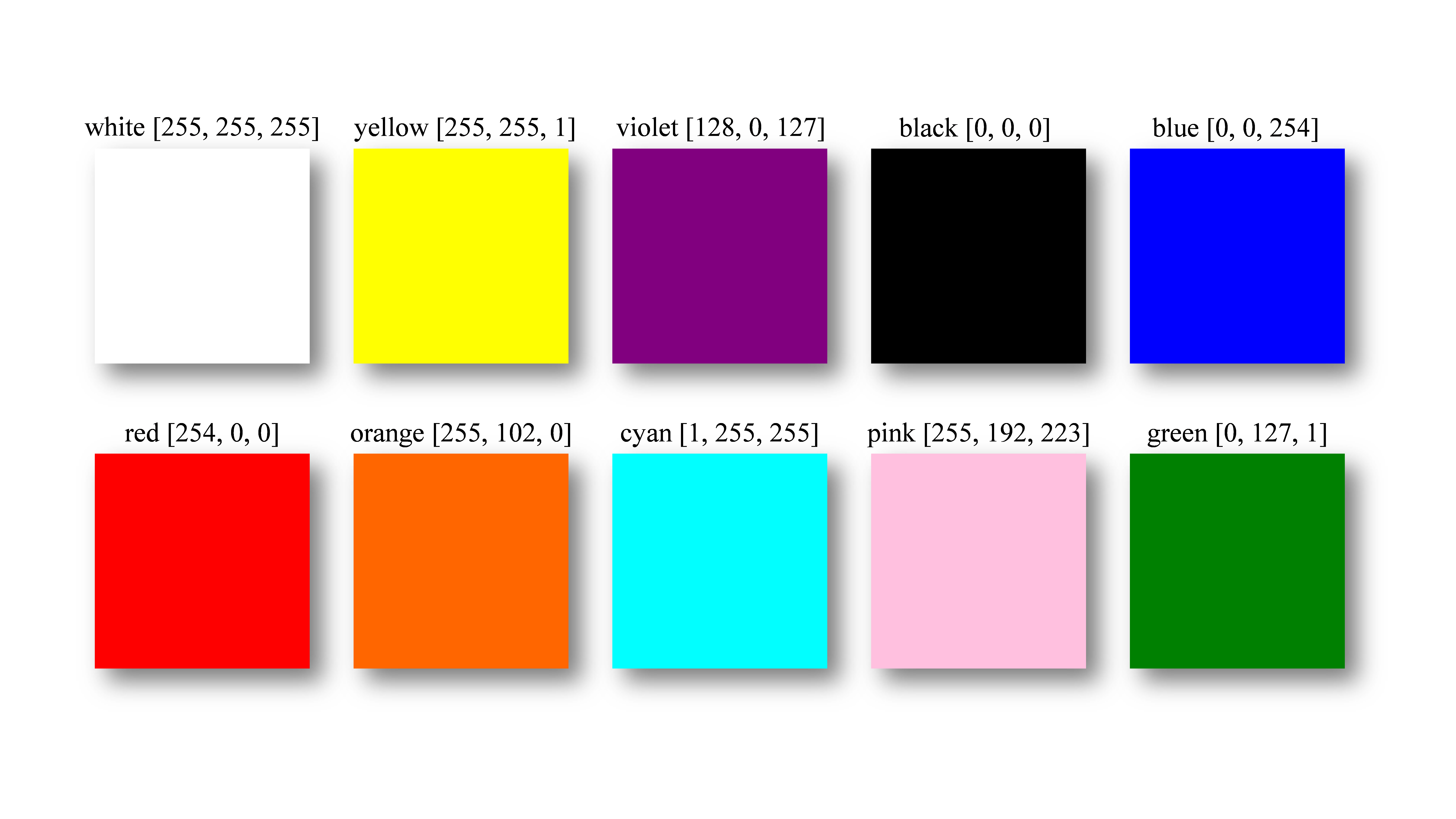

**Figure S2** Ten distinguishable colors of all squares used in behavioral change detection task.

**Appendix S3. Correlations between the severity of ADHD symptoms and MMN**

As an exploratory analysis, we assessed whether individual differences in the severity of ADHD symptoms were linked to susceptibility to task-irrelevant auditory distractors, and performed partial correlation analyses between ADHD symptom scores and auditory MMN amplitudes in children with ADHD, controlling for age as a covariate. The results (Figure S3) revealed that in both experiments, auditory MMN amplitudes were not significantly correlated with any of the symptom scores, including inattentive, hyperactive/impulsive, or total scores (*p*s > .528, uncorrected).

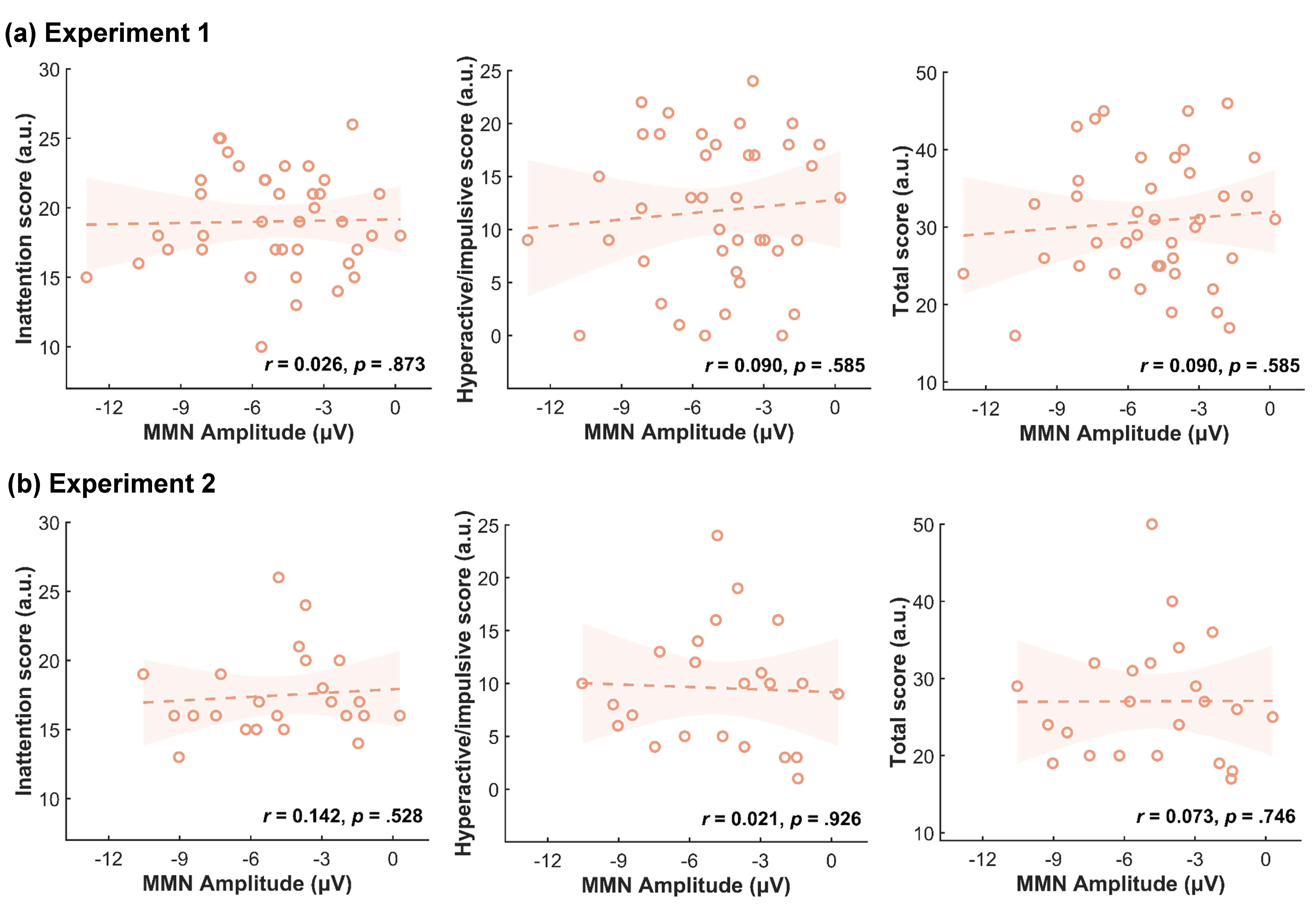

**Figure S3** Correlation between MMN amplitudes and inattentive scores, hyperactive/impulsive scores, or total scores for children with ADHD in (a) Experiment 1 and (b) Experiment 2. Shading represents the 95% CIs of the trend line.

**Appendix S4. Comparisons of visual WM capacity across Experiments 1 and 2**

To compare group differences in the visual WM capacity between Experiments 1 and 2, a two-way ANCOVA with age as a covariate was performed on the K-scores. The analysis revealed a significant main effect of group (*F*_(1,141)_ = 5.613, *p* = .019,$\eta_{p}^{2}$ = 0.038), suggesting that children with ADHD exhibited lower visual WM capacity than TD children did. In contrast, no significant main effect of experiment (*F*_(1,141)_ = 0.029, *p* = .865,$\eta_{p}^{2}$ < 0.001) or interaction between experiment and group (*F*_(1,141)_ = 0.188, *p* = .665,$\eta_{p}^{2}$ = 0.001) was found.

**Appendix S5. Comparisons of MMN amplitudes across Experiments 1 and 2**

To examine whether increasing audiovisual competition altered the neural responses to auditory distraction in the two groups of children, we also performed a one-way ANCOVA using age as a covariate on the MMN amplitudes from both experiments combined. The results revealed a significant main effect of experiment (*F*_(1,141)_ = 6.611, *p* = .011,$\eta_{p}^{2}$ = 0.045), along with a significant interaction between experiment and group (*F*_(1,141)_ = 4.095, *p* = .045,$\eta_{p}^{2}$ = 0.028). Post hoc analysis indicated that the MMN amplitudes of the TD group were significantly greater in Experiment 1 than in Experiment 2 (*p* = 0.001), whereas no significant difference was found in the MMN amplitude of the ADHD group between the two experiments (*p* = 0.720).
